# Quantifying maternal antibody transfer to colostrum and cord blood reveals virus-specific selectivity in dogs

**DOI:** 10.1101/2025.11.24.690260

**Authors:** Serena Teh, Lotta H Truyen, Sarah Woodyear, Jessica Palmeri, Ana Alice Pimenta-Pereira, Alexis AE Peprah, Ella Lanman, Tom M Lonergan, Abigail Reid, Soon Hon Cheong, Sarah L Caddy

## Abstract

**Background:** Neonatal infections are a leading cause of mortality in dogs, with up to 30% of puppies dying within the first three weeks of life. During this period of immune development, protection is highly dependent on maternal antibodies (MatAbs) transferred across the placenta and via colostrum. Despite the critical importance of this transfer, little is known about the biological or clinical factors that determine its magnitude, whether specific antibodies are preferentially transferred, or how these processes vary across a broad population of dogs.

**Methods:** To quantify and explore the determinants of MatAb transfer in dogs, we analyzed matched maternal serum, cord blood, and colostrum samples collected from 44 client-owned dams undergoing cesarean section at a university veterinary hospital. Total IgG and virus-specific antibodies against canine parvovirus (CPV) and canine distemper virus (CDV) were analyzed. We also evaluated the influence of maternal factors, including age, breed, body weight, parity and litter size on MatAb transfer efficiency.

**Results:** Across this diverse population, we observed limited transplacental transfer of MatAbs (4.5-6% maternal titer), in agreement with previous studies and as expected with the endotheliochorial placenta of dogs. In contrast, virus-specific IgG was highly enriched in colostrum, with 10.7-fold (CPV) and 8.1-fold (CDV) increases relative to serum. Transfer efficiency was significantly greater for virus-specific antibodies than for total IgG (3.2-fold), suggesting selective enrichment of antiviral antibodies during colostrogenesis. Maternal serum antibody titer emerged as the primary factor influencing the efficiency of antibody transfer.

**Conclusions:** These findings provide the most comprehensive quantification to date of MatAb transfer routes in dogs, revealing preferential transfer of virus-specific IgG to colostrum and highlighting the crucial role of colostrum intake for neonatal immunity. This work establishes a foundation for identifying antibody characteristics that influence MatAb transfer efficiency and reiterates the importance of ensuring dams have adequate titers of virus-specific IgG prior to breeding.

## INTRODUCTION

Neonatal infections are a leading cause of puppy mortality, with young animals especially vulnerable whilst their immune systems are still developing. Up to 30% puppies born die within the first 3 weeks of life, and a proportion of these are caused by early infections (1,2). To support this period of immunodeficiency, maternal antibodies (MatAbs), sometimes referred to as maternally-derived antibodies, are delivered to pups from the mother across the placenta and in colostrum. It has been shown that if inadequate antibody transfer occurs, puppy mortality is significantly higher (1).

The dominant route of MatAb transfer varies across mammalian species. In humans the majority of maternal IgG received by infants is transferred across the placenta. In contrast, in ruminants and horses all maternal IgG is transferred via colostrum. Dogs are an intermediate species; it has previously been estimated that 5-10% total maternal IgG crosses the placenta (3,4). This is understood to be due to the different placental structure of the different species; three tissue layers separate the maternal and fetal circulations in the human hemochorial placenta, whereas four tissue layers are present in the canine endotheliochorial placenta and six layers in the epitheliochorial placenta of ruminants and horses (5). The majority of MatAb transfer in dogs is therefore via the colostrum (6,7). However, previous studies quantifying MatAb transfer routes have had their limitations; most have only studied a small number of research dogs (4,8,9), and only quantified total IgG or IgG specific for a single pathogen.

In this study we sought to quantify MatAb transfer across the placenta and via colostrum in a diverse group of client-owned dogs. To obtain samples from dogs at birth we recruited cases undergoing cesarian sections at a University Hospital. This approach enabled us to collect cord blood from puppies, which represented maternal IgG transferred across the placenta, as well as the first colostrum from the dam. Matched serum from each dam was also collected. We comprehensively analyzed samples from this unique cohort for MatAbs specific for two viruses to which neonates are highly susceptible; canine parvovirus (CPV) and canine distemper virus (CDV) (10,11). These results were compared with total IgG titers and revealed an unexpected preferential transfer of anti-viral antibodies compared to total IgG. Overall this analysis has allowed us to precisely determine the extent to which MatAbs are transferred across the placenta and mammary gland in dogs, explored the factors impacting canine MatAb transfer, and reinforced the importance of colostrum ingestion for puppies.

## METHODS

### Sample collection

To study MatAbs, samples were collected from 44 client-owned dogs undergoing cesarean sections at Cornell University Hospital for Animals (CUHA, IACUC approval 2023-0003). Maternal blood was collected from either the cephalic or saphenous vein. Umbilical cord blood samples were taken immediately after surgical delivery of pups as follows; the umbilical cord was clamped at either end to detach the placenta from the puppy. Whenever possible, a syringe and needle were used to aspirate the blood from the cord. If this method was not feasible, the clamp closer to the puppy was removed and a 1.5 ml microcentrifuge tube used to collect cord blood manually expressed from the blood vessels. Samples of both live births and stillborns were collected and used in analysis. Colostrum was milked out of at least 4 teats and then pooled and stored in a single 1.5ml microcentrifuge tube (variation has been reported between teats (7)). All samples were stored at 4°C for up to 24 hours until further processing, whereby maternal and cord blood samples were centrifuged for 5 minutes at 8000 rpm to separate serum. All samples were stored long term at -80°C. Basic clinical and demographic data for each dam were recorded, including age, breed, weight, body condition score, parity, litter size and any known vaccination history.

### Production of canine parvovirus (CPV) virus-like particles (VLPs)

For the production of CPV VLPs, the Bac-to-Bac Baculovirus Expression system was used in DH10 *E. coli* bacteria. Plasmid coding for the CPV VP2-capsid protein was purified using a commercial kit (Monarch Plasmid Miniprep kit) according to manufacturer recommendations. Sf9 insect cells were transfected with VP2 BacMid using Cellfectin-II transfection reagent (Invitrogen), using 8μl Cellfectin-II and 1μg BacMid in 200μl per T-25 flask with serum-free Grace Media. After incubating for 4 hours at 27°C, the transfection mix was removed, and Grace Media with 10% fetal calf serum (FCS) was added and incubated for 4-7 days at 27°C. To harvest the P0 virus stock, the media from the cell culture was collected and then centrifuged for 5 min at 500g. Then 200μl of P0 viral stock was added and incubated for 4-5 days in a humidified incubator at 27°C. To amplify the P1 virus in Sf9 cells, supernatant was collected from transfected Sf9 cells and centrifuged for 5 min at 500g, and 100μl supernatant transferred to previously seeded Sf9 insect cells. Cells were incubated for 4-5 days in a humidified incubator at 27°C then virus harvested. Protein production was subsequently performed in Hi5 cells infected with P2 virus under the same conditions. VLPs were purified using a cesium chloride (CsCl) gradient, followed by dialysis to remove residual CsCl. VLP production was confirmed by SDS-PAGE with Coomassie staining and western blotting. Protein concentrations were determined using both a bicinchoninic acid (BCA) assay and a Qubit fluorometric assay.

### Canine distemper virus (CDV) propagation

Vero cells stably expressing canine SLAM (CD150) were incubated at 37°C and 5% CO_2_ in DMEM with 50 I.U./mL penicillin, 50μg/mL Streptomycin and 5% FCS (complete media). Cells were infected with CDV (Onderstepoort strain, provided by Diego Diel, Cornell University) at an MOI of 0.01 in 5 ml of infection media with 2% FCS. The virus-media mix was incubated with cells for one hour at 37°C with gentle rocking side to side every 15 minutes. Complete media was then added and the flask was incubated and checked daily for cytopathic effect (CPE). Once the CPE occurred, supernatant was collected, centrifuged at 1600rpm for 5 min at 4°C, then aliquoted and stored at -80°C. A TCID50 assay was used to determine viral titer. A total of 2 x 10^4 Vero-dogSLAM cells were seeded per well of a 96-well plate in a total volume of 50μl. A 10-fold dilution of supernatant containing virus was diluted in complete media, and then 50μl per well of each dilution was added in quadruplicates to the seeded cells. The last row of wells remained uninfected as a negative control. The plate was incubated until a clear CPE was visible and the titer was determined by TCID50 using the Reed-Muench method.

### ELISAs

For virus-specific ELISAs, high binding ELISA plates (Greiner) were coated with either 15ng of CPV-VLPs in PBS per well, or 1×10^5^ TCID50 CDV particles. Plates were incubated overnight at 4°C, then washed with PBS-0.1% Tween 20 (‘PBS-T’) and blocked with 5% milk in PBS-T. After washing with PBS-T, serially diluted dam serum (1:800 starting dilution), cord serum (1:100 starting dilution), or colostrum (1:1600 starting dilution) was added. Samples were incubated for 2 hours at 37°C. Following washing with PBS-T, anti-canine IgG (H+L) conjugated to horse radish peroxidase (HRP) (Invitrogen, CAT#PA1-29738) or anti-canine IgG2 (Bethyl Laboratories CAT#A40-121P) was added for 1 hour. After a final wash step TMB substrate (Abcam) was added for 10 minutes. This reaction was stopped using 1M H_2_SO_4_ and absorbance measured at OD450 (BioTek Cytation 7 Cell Imaging Multimode Reader, Agilent). Each sample was tested in duplicate, and a positive sample (pooled serum samples from 5 healthy, vaccinated dogs) as well as a virus-specific IgG negative control (Marshall Bioresources, NY) were included on every plate. To correct for background signal, the optical density (OD) values from wells coated with PBS only were subtracted from the OD values of wells coated with VLPs or virus. Corrected OD450 values were plotted against dilution in Graphpad Prism, and non-linear regression analysis with interpolation was applied to calculate endpoint titers. The endpoint titer was defined as the reciprocal of the dilution at which OD450 crossed the detection threshold, calculated as the mean OD450 of all PBS-coated wells plus three standard deviations. Results were only accepted if the non-linear regression fit for both the standard curve and the sample curve had an R^2^ ≥ 0.9.

Indirect ELISA assays were developed in house to quantify total IgG in clinical samples. Rabbit anti-canine IgG (Invitrogen, CAT#SA5-10309) was coated on ELISA plates at 1μg/ml concentration in PBS overnight at 4°C. The wells were washed 3 times with PBS-T, then blocked with 5% milk in PBS-T. After washing with PBS-T, serially diluted dam serum, cord serum or colostrum was added. Each sample was tested in duplicate. For the standard curve, serially diluted canine IgG (Southern Biotech, CAT#0129-01) starting at 5μg/ml was added in duplicate to the plate. Samples were incubated for 2 hours at 37°C. After washing, the TMB substrate was added and 1M H_2_SO_4_ was used to stop the reaction. Absorbance was measured at OD450. Background signal was corrected for by subtracting PBS only values as with virus-specific ELISAs. Corrected OD450 values were plotted against dilution factor in GraphPad Prism, and non-linear regression analysis with interpolation to standard curve was performed to calculate the IgG concentration in each sample. Results were only accepted if the non-linear regression fit for both the standard curve and sample curve had an R^2^ ≥ 0.9.

### Statistical Analysis

Based on preliminary data and paired sample comparisons, recruitment of 25 dogs was estimated to provide 80% power (α = 0.05) to detect a standardized difference of 0.6 in antibody transfer ratios between maternal and neonatal samples, a magnitude similar to that observed in human studies (12,13). To ensure robust analysis across a naturally occurring cohort, we ultimately collected matched samples from 44 dams, exceeding the minimum requirement. Puppies from the same litter were considered biological replicates. All data analysis was performed using GraphPad Prism (version 10.3.1). End-point titer outcomes of CPV and CDV-specific IgG were non-normally distributed, as were total IgG concentrations, and so all were analyzed by Kruskal-Wallis test and Dunn’s multiple comparisons test. Transfer ratios between dam and colostrum IgG as well as dam and cord IgG were analyzed by Mann-Whitney tests. Correlation analysis included nonlinear regression, Pearson’s correlation for normally distributed variables, and Spearman’s rank correlation for non-normally distributed variables. Statistical differences were considered significant where *p* ≤ 0.05 for all comparisons.

## RESULTS

### Study population

A total of 44 female dogs (‘dams’) that presented at Cornell University Hospital for Animals for caesarean sections were enrolled in this study. The dams were an average 3.6 years old (range 1-8 years), and multiple breeds were represented including brachycephalic breeds (American Bulldog (4), French Bulldog (8), English Bulldog (2), Bully cross (2), Pitbull (1), Chihuahua (1), Shih Tzu cross (1)) and non-brachycephalic breeds (Labrador (4), Corgi (3), Golden Retriever (3), Leonberger (2), others (12)). All caesarean sections were emergency procedures except for a single elective procedure. Demographic and clinical details for each case are presented in Table 1.

**Table 1.** Maternal demographic and clinical details of the study cohort.

| Table 1. Maternal demographic and clinical details of the study cohort |  |  |  |  |  |
| --- | --- | --- | --- | --- | --- |
|  | Range | Mean | Standard deviation | Factor | Cases |
| Age of dam (years) | 1-8 | 3.6 | 1.8 | <2y | 4 (9%) |
|  |  |  |  | 2-3y | 18 (41%) |
|  |  |  |  | 4-5y | 15 (34%) |
|  |  |  |  | 6y+ | 7 (16%) |
| Litter size | 1-11 | 6.5 | 2.6 | <3 | 1 (2%) |
|  |  |  |  | 3-6 | 24 (55%) |
|  |  |  |  | >6 | 19 (43%) |
| Parity | 1-4 | 1.6 | 0.9 | Primiparous | 19 (43%) |
|  |  |  |  | Multiparous | 23 (52%) |
|  |  |  |  | Unknown | 2 (5%) |
| Breed |  |  |  | Brachycephalic | 19 (43%) |
|  |  |  |  | Non-brachycephalic | 25 (57%) |
| Body Condition Score (1-5) | 1-5 | 3.3 | 0.6 | ≤2 | 2 (5%) |
|  |  |  |  | 2.5-3.5 | 30 (68%) |
|  |  |  |  | ≥4 | 9 (20%) |
|  |  |  |  | Not recorded | 3 (7%) |
| Vaccination status |  |  |  | Up to date | 20 (45%) |
|  |  |  |  | Not up to date | 6 (14%) |
|  |  |  |  | Unknown | 18 (41%) |

### Transfer of canine virus-specific maternal antibody

IgG is the major antibody isotype transferred from mother to infant in all mammalian species (14,15). For puppies, transfer of MatAbs specific for life-threatening pathogens is of critical importance for neonatal health, and so we first wanted to quantify IgG for two significant canine pathogens; canine parvovirus (CPV) and canine distemper virus (CDV). We also reasoned that the majority of dams would be seropositive to these viruses due to core vaccine recommendations (16), thus studying transfer of these MatAbs would be readily achievable. We quantified virus-specific IgG in dam serum and compared this to MatAbs detectable in colostrum. We also compared virus-specific IgG in dam serum with that detected in serum separated from cord blood, making this the first reported use of cord blood sampling to quantify MatAbs delivered transplacentally in dogs. Our cord sampling strategy enabled precise measurement of placental antibody transfer, overcoming the limitations of prior methods that have relied on blood sampling from newborn puppies. Direct sampling of puppies is more ethically sensitive, technically challenging, and potentially confounded by early colostrum intake.

Virus-specific IgG titers for all samples in our cohort were quantified using in-house ELISAs. Serial dilutions of each sample were generated, and the endpoint titer was defined as the highest dilution at which antibody binding to the viral antigen remained detectable. Endpoint titers are presented as the reciprocal of this final positive dilution. A small number of dams had undetectable titers of IgG specific for CPV (3 dams, 6.8%) or CDV (6 dams, 13.6%), with 2 of these dams having no detectable IgG to both viruses (4.5% dams). The presence of seronegative individuals in our breeding cohort was a concern, but these rates were lower than those reported in a previous US study of hospitalized dogs, which found 19% and 50% seronegativity for CPV and CDV, respectively (17). We omitted colostrum or cord samples from cases with undetectable virus-specific IgG from further study. Additionally, in a small proportion of cases it was not possible to collect the full clinical dataset (dam serum, colostrum, cord blood). The number of samples studied for each virus-specific IgG are listed in Table 2.

**Table 2.** Virus-specific IgG quantified by ELISA. Endpoint titers are presented as the reciprocal dilution at which antibody binding to antigen was detectable. Transfer ratios were calculated as the IgG endpoint titer in colostrum or cord divided by the corresponding IgG titer in the matched dam serum.

| Virus |  | IgG endpoint titer |  |  | Transfer ratio |  |
| --- | --- | --- | --- | --- | --- | --- |
|  |  | Dam | Colostrum | Cord | To colostrum | To cord |
| Canine parvovirus (CPV) | Mean | 47474 | 404381 | 1909 | 10.7 | 0.045 |
|  | Standard deviation | 54254 | 515169 | 3070 | 8.0 | 0.057 |
|  | Number of samples analyzed | 41 | 40 | 124 | 37 | 118 |
| Canine distemper virus (CDV) | Mean | 8968 | 49991 | 391 | 8.1 | 0.060 |
|  | Standard deviation | 21533 | 83311 | 558 | 6.3 | 0.047 |
|  | Number of samples analyzed | 38 | 34 | 54 | 35 | 50 |

IgG titers for each dam with detectable virus-specific IgG are presented in Figure 1A and 1B, where each point represents a single dam and her litter. The average (mean) cord IgG titer within each litter is shown as a single point. We found that virus-specific IgG in colostrum was significantly higher (p=0.0003 and p=0.0029 for CPV and CDV respectively, determined by Kruskal-Wallis test) than IgG circulating in dams for both viruses, as summarized in Table 2. In contrast, the average cord IgG titer for each litter was significantly lower than that detected in corresponding dam serum. To evaluate how MatAb transfer varied at the individual level we calculated transfer ratios for every colostrum and cord sample for which the titer was above the lower limit of quantification. The results are presented in Figure 1C. This provides clear evidence that IgG is actively concentrated into the colostrum, resulting in IgG levels that are 8.1-fold higher for CDV and 10.7-fold higher for CPV compared with those in the dam’s serum. Cord sample analysis demonstrated that approximately 5% virus-specific maternal IgG is able to cross the placenta.

**Figure 1.**
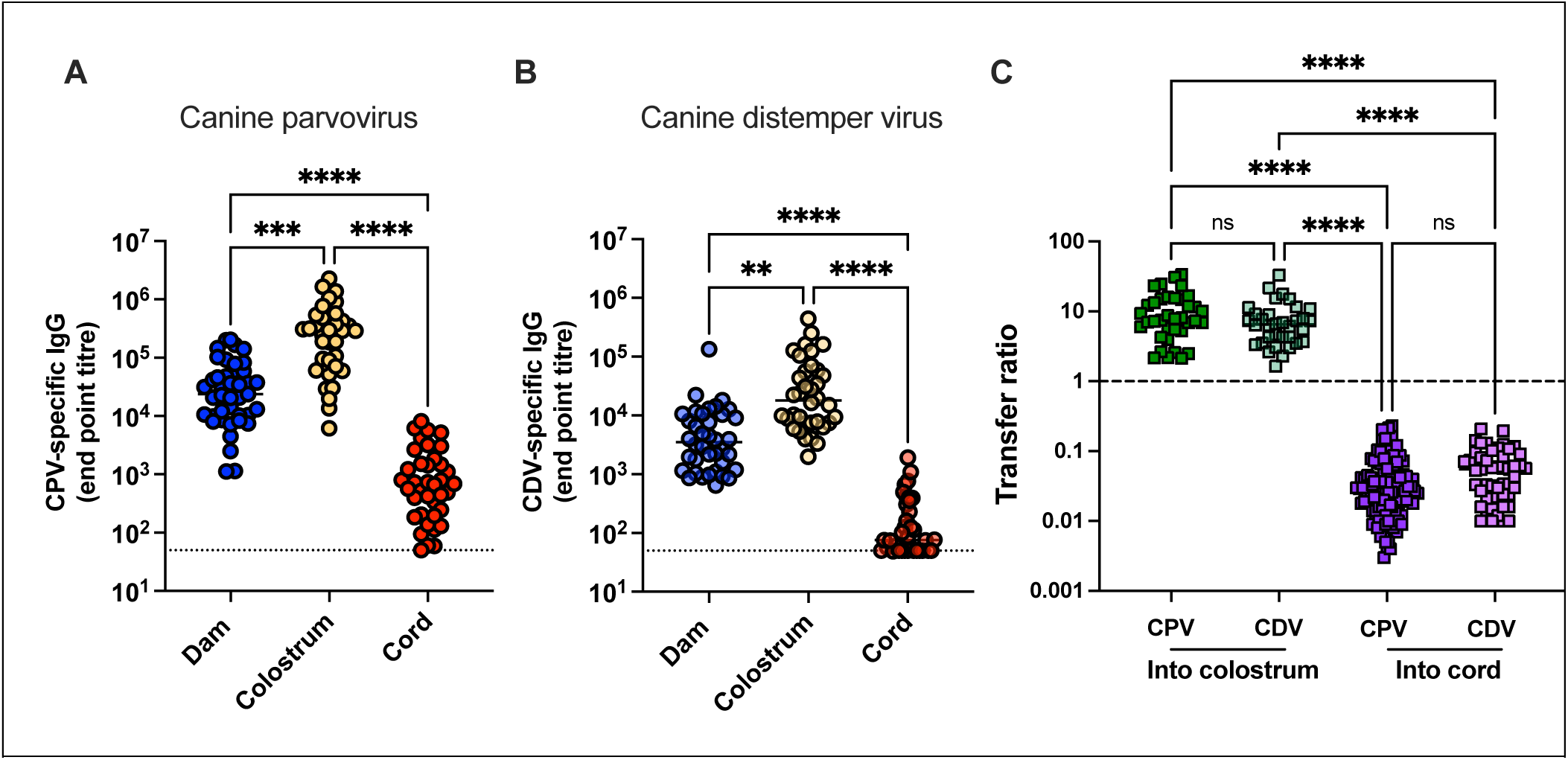
Virus-specific IgG quantified in dam serum, colostrum and cord samples. End point titers of IgG specific for A) canine parvovirus (CPV) and B) canine distemper virus (CDV) were quantified in clinical samples collected from dams and umbilical cords during cesarean sections. Transfer ratios of IgG between dam serum and colostrum/cord for puppies are shown in C. Dotted lines in A/B represent lower limit of quantification. Dashed line in C represents a 100% transfer efficiency (equivalent IgG levels between samples). Significance was assessed using Kruskal Wallis tests with multiple comparisons. Asterisks denote statistical significance: *p* <0.01 (**)*, p* <0.001 (***), and *p* <0.0001 (****).

To extend this analysis, we performed CPV-based ELISAs to quantify different MatAb isotypes and subclasses. Analysis of a subset of 6 groups of samples (dam, colostrum, cord) confirmed that IgM and IgA titers in all samples were very low compared to the IgG isotype, so further quantification was not performed (data not shown). In several species, certain subclasses of IgG are preferentially transferred to neonates (18,19). There is only limited analysis of the roles of different IgG subclasses in dogs at any life stage, and no previous work has evaluated transfer rates of maternal IgG subclasses. To begin evaluating transfer differences, we modified our in-house indirect CPV-specific IgG ELISA to quantify CPV-specific IgG2 antibodies. IgG2 has been reclassified as a combination of IgG subclass B, and IgG subclass C (20), but commercially available secondary antibodies are unable to distinguish between these subsets (21). We quantified CPV-specific IgG2 MatAbs in a randomly selected subset of samples (29 dams, with matched colostrum and cords) and demonstrated that there was no preferential transfer of CPV-specific IgG2 relative to all IgGs in our cohort (Supplementary Figure 1).

### Evaluation of total IgG in dam serum, colostrum and cord samples

Previous studies of MatAb transfer have largely focused on total IgG transferred, and not virus-specific MatAbs (7,9,22,23). We therefore sought to include total IgG quantification in our study, to compare with both previous reports and with our virus-specific MatAb analysis. IgG titers from 24 groups in the cohort (selected for having adequate remaining sample volume) are shown in Figure 2A, with each point representing the IgG titer of one dam or one colostrum sample, or the average cord IgG titer within one litter. The mean titers of total IgG in our cohort of dam serum, colostrum and cord were quantified as 7.9mg/ml, 19.5mg/ml and 0.59mg/ml respectively. Our findings were in close agreement with previously reported results which quantified the average total IgG in dam circulation as 8.1mg/ml and in colostrum to be 19.3-20.8mg/ml (7,23). We also compared our cord IgG titers with IgG titers quantified from serum drawn from puppies at birth, which also show similar results of 0.3mg/ml (24,25) and 1.2mg/ml (22). This confirms collection of umbilical cord blood as an alternate, non-invasive strategy to collect serum from puppies to determine the efficiency of placental transfer of IgG.

**Figure 2.**
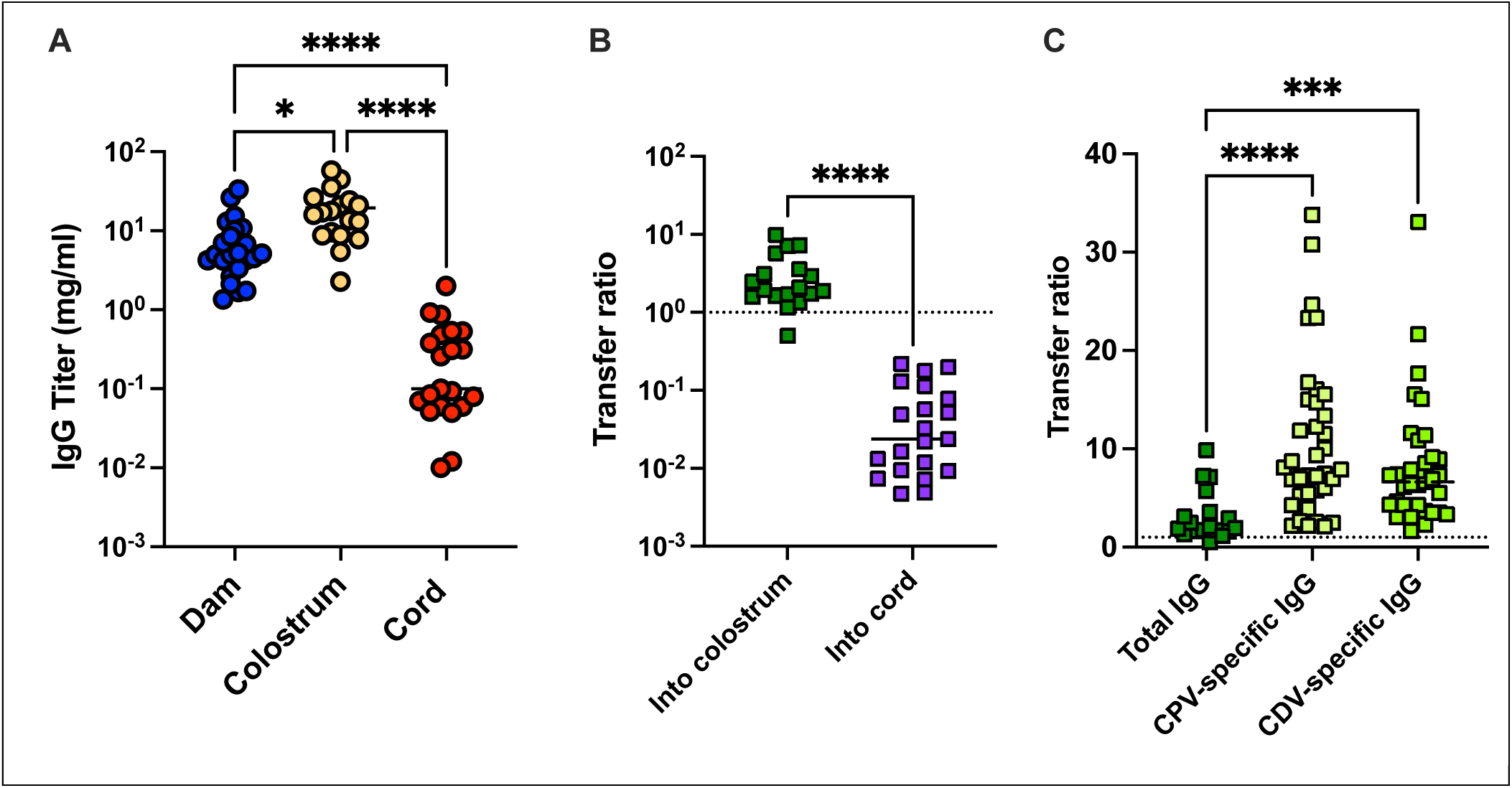
Total IgG titers quantified in dam serum, colostrum, and cord samples. (A) Total IgG titers were quantified in clinical samples by indirect ELISA; dam n=24, colostrum n=18, cord n=21 litters. (B) Transfer ratios of IgG via colostrum (n=18) or cord (n=21) calculated from total IgG titers. The dotted line represents transfer ratio of 1, representing an equivalent concentration of IgG between samples. (C) Transfer ratio of total IgG compared to virus-specific IgG from maternal serum into colostrum. Significance was assessed by a Kruskal Wallis test with post-hoc multiple comparisons (A, C) and a Mann Whitney test (B). Asterisks denote statistical significance: *p* <0.05 (*)*, p* <0.001 (***) and *p* <0.0001 (****).

Next, we calculated the transfer ratio of total IgG via colostrum and placenta as shown in Figure 2B. Similar to virus-specific IgG, our results support the conclusion that puppies receive the majority of maternally derived IgG through ingesting colostrum. Unexpectedly however, the concentration of total IgG in colostrum relative to dam sera is significantly lower than the concentration of virus-specific IgG in colostrum relative to dam sera as shown in Figure 2C. This data suggests that virus-specific IgG is being selectively transported into the colostrum for transfer to the puppies. In contrast, the transfer ratios for total IgG and virus-specific IgG from dam sera to cord are highly comparable (total IgG 0.059, CPV-specific IgG 0.045 and CDV-specific IgG 0.060).

### Correlation between maternal antibody titers in dam, colostrum and cord samples

Given the substantial variation in antibody titers in colostrum and cord samples relative to dam serum, we sought to identify which factors were important for canine MatAb transfer. We began by evaluating the relationship between IgG titers in the dam serum, and titers in colostrum and cord samples. Whilst studies in humans, sheep and pigs have identified a positive correlation between maternal IgG titer and colostrum (26–28), prior studies in dogs have not identified an association (7). In addition, multiple studies in humans have demonstrated a strong correlation between maternal and cord IgG titers, especially for virus-specific antibodies (29–31), but this has not previously been explored in dogs.

The correlations between dam serum IgG titers and cord or colostrum IgG titers for our two representative viruses are presented as scatterplots in Figure 3. Non-linear regression analysis identified a consistent positive trend across all virus-specific MatAb curves. Due to non-parametric data distribution, correlations were assessed using Spearmans rank correlation (π). This demonstrated highly significant associations for each sample type and virus (*p* < 0.0001).

**Figure 3.**
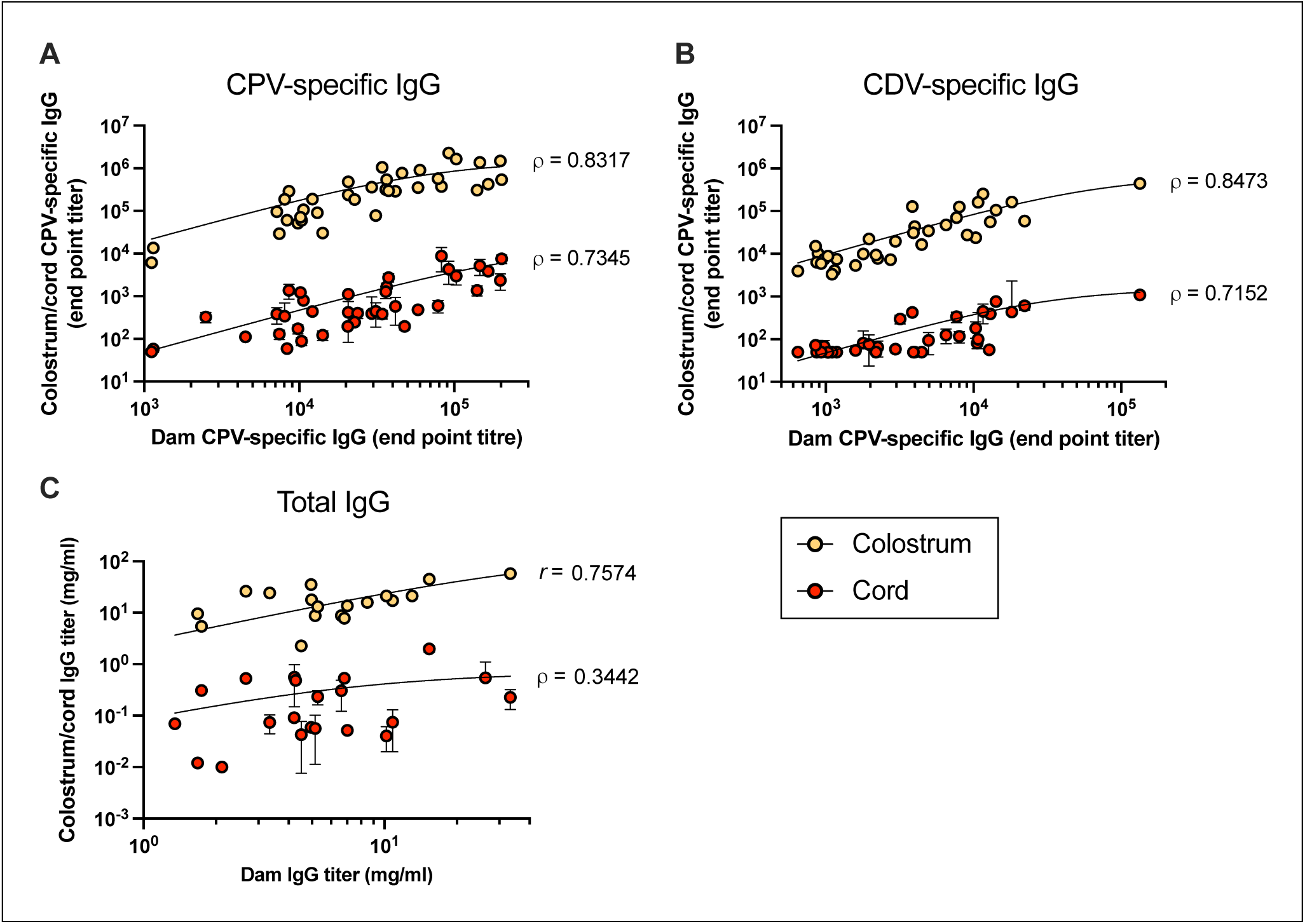
Correlation between dam IgG titer and IgG titers in colostrum and cord samples. Scatterplots of IgG titers for A) CPV (colostrum n=36, cord n=38 litters) and B) CDV (colostrum n=35, cord n=35 litters), and C) total IgG (colostrum n=18, cord n=22 litters) as determined by ELISA. Non-linear regression curves were fitted to the data. Statistical significance was assessed using Pearson’s correlation coefficient (*r*) for normally distributed data (total IgG colostrum) and Spearman’s rank correlation coefficient (*ρ*) for non-parametric data.

The strong association between dam and colostrum titers was unexpectedly different from results by Mila et al (7). However, in this earlier study, total IgG and not virus-specific IgG was quantified in clinical samples. To determine if antibody specificity affected this relationship we assessed the correlation between total IgG titers in each sample as shown in Figure 3C. We found a moderate positive correlation between dam and colostrum total IgG titers (data normally distributed, Pearson’s correlation *r*=0.7574, *p*=0.0003), and no significant correlation between dam and cord total IgG titers (data non-normally distributed, Spearman’s correlation *r*=0.3442, *p*=0.1266).

### Correlation between MatAb transfer and maternal variables

To identify any demographic or clinical factors that correlated with MatAb transfer, we analyzed how IgG titers in each sample varied with parity, age of dam, weight of dam and litter size. Figure 4 presents the results of CPV-specific IgG analysis and comparable results were obtained with CDV-specific IgG analysis (data not shown). Spearman’s rank correlation was performed for each variable, and no correlation between virus-specific IgG and any of these variables was identified (*p*>0.05). Vaccine status was also evaluated, but no differences between dogs confirmed to be up to date with vaccines (20 dogs) and those that were not up to date (6 dogs) were evident (data not shown). We also performed the same analysis with total IgG results (supplementary Figure 2), and again did not observe any correlations between MatAbs and the factors studied.

**Figure 4.**
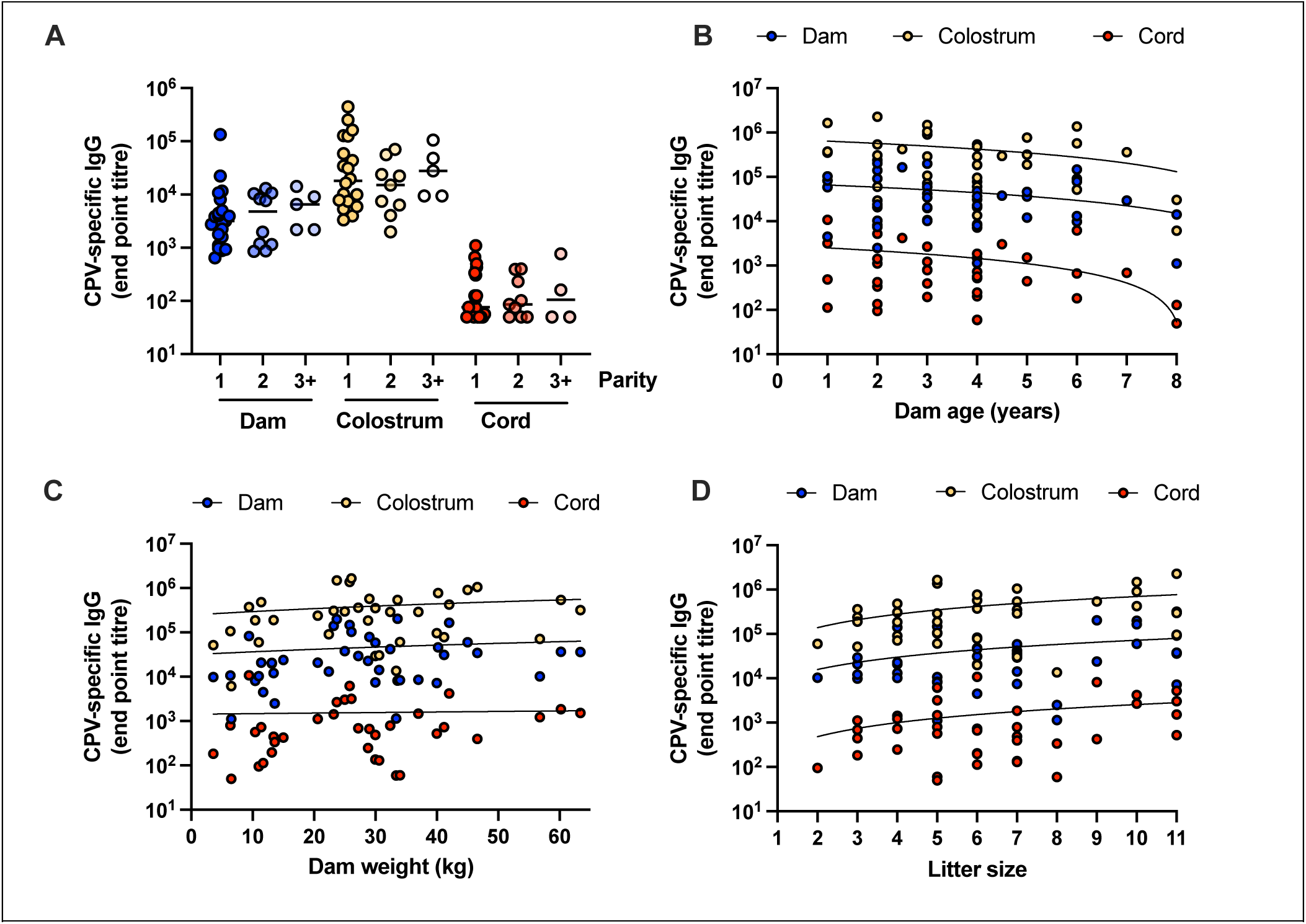
Correlation between CPV-specific IgG titers and biological variables. All values were determined by CPV-specific IgG ELISA. (A) End point titers of CPV-specific IgG from dam, colostrum and cord plotted against parity. Horizontal bar represents mean titer of each group. (B-D) End point titers of CPV-specific IgG plotted against dam age, dam weight and litter size respectively. Non-linear regression curves were fitted to the data.

### Heterogeneity in cord IgG titers and early protection from viral infection

This study is the first to report analysis of MatAb titers in canine cord blood samples. As shown in Figure 1, we demonstrate that approximately 5% of circulating virus-specific IgG in the dam is transferred across the placenta. However, we observed significant variation across litters and wanted to explore if there was also variation in IgG titer *within* litters. Figure 5 presents the CPV-specific cord IgG titers of 122 puppies from 38 litters. Of the 6 litters for which no cord results were available, 3 of these were from dams with no CPV-specific IgG, and the remainder had insufficient sample available or sample was not collected. Though the majority of litters have cord IgG titers that cluster closely together, there are 9 litters with a standard deviation >1000.

**Figure 5.**
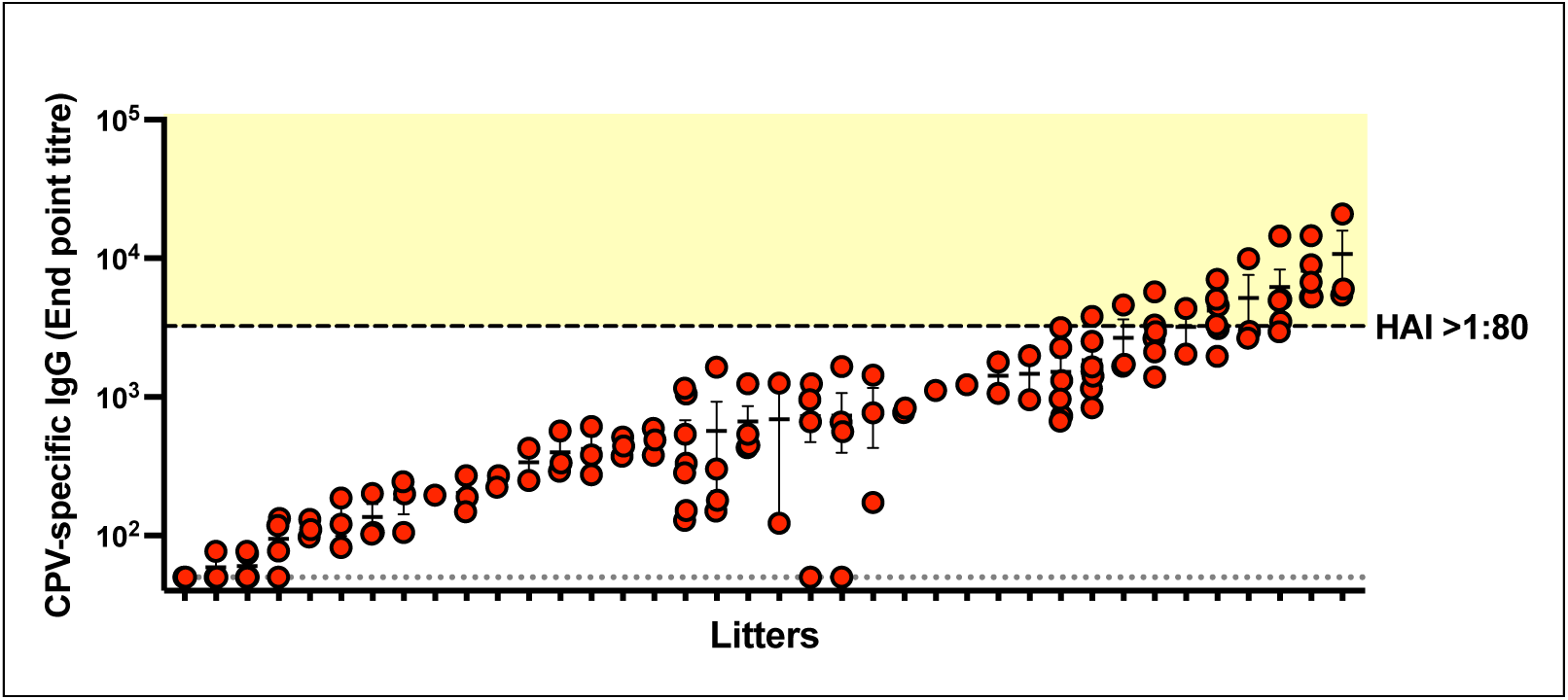
Titers of CPV-specific IgG in cord samples within each litter. Cord samples from each litter (n=38 litters, 1 – 7 cords per litter) are plotted together, arbitrarily ordered according to mean titer. Error bars represent standard error of the mean. Dotted line represents lower level of quantification of ELISA assay. Dashed line corresponds to an ELISA end point titer of 3239, and hemagglutination inhibition titer (HAI) >1:80. Shaded yellow area represents protective IgG titers.

It was apparent that a proportion of puppies were born with moderately high levels of CPV-specific IgG. We asked if this level would be consistent with protection at birth. Previous studies have reported a hemagglutination inhibition (HAI) titer of >1:80 corresponds to protection (4), and that HAI and ELISA titers are highly correlated (32). Our standard positive control was submitted to the Cornell Animal Health Diagnostic Centre for quantification by HAI. This established that an HAI titer of 1:80 corresponded to an end point titer of 3239 in our in-house ELISA. This is highlighted in Figure 5 as a dashed line, with samples >1:80 falling within the shaded yellow area. A total of 22 (17.2%) puppies were born with sufficient IgG to render them protected from CPV at birth without ingesting any colostrum.

## DISCUSSION

This study presents a comprehensive analysis of MatAb transfer, both transplacental and via colostrum, in a varied population of client-owned dogs. Whilst colostral MatAb transfer has been carefully evaluated previously (7), quantification of transplacental transfer has been limited to small groups of puppies that were blood sampled as neonates (22,24,33). To expand on this understanding, we now provide the first report of placental transfer evaluation by collection of umbilical cord blood samples. This is an approach widely used in human studies (12,13) but has not previously been reported in dogs. Blood sampling from puppies prior to their first suckle limits the number of samples collected due to the invasive nature of this method and reduced likelihood of owner consent when testing client-owned animals. We found that quantification of total IgG in cord samples showed very similar results to total IgG titers in pup serum collected before colostrum ingestion; 0.46mg/ml in cord serum, compared to previous reports of 0.3-1.2 mg/ml in pup serum (22,24,33). Furthermore, we demonstrated that virus-specific cord IgG titers were 4.5% (CPV-specific) and 6.0% (CDV-specific) of dam IgG titers, remarkedly comparable to the 5.7% reported by Pollock and Carmichael, despite this original study being conducted in research kennels and our cohort being client-owned dogs (4). This confirms that collection of cord blood is a suitable method for studying placental transfer of canine MatAbs, and also verifies that placental transfer only contributes a small proportion of MatAb delivery to pups.

Using our valuable cohort of samples, we quantified virus-specific IgG for two important canine viruses (CPV and CDV), as well as total IgG. Evaluation of canine MatAbs using both approaches has not previously been conducted on the same sample set. Interestingly, we observed greater transfer rates of virus-specific IgG into colostrum (8.1-fold for CDV, 10.7-fold for CPV) than for total IgG (3.2-fold). This provides evidence for selective transfer of MatAbs, which is the first report of this process in dogs. Identification and exploration of this phenomenon in other domestic species is also limited. A single report in cows has provided evidence of selective-transfer of pathogen-specific IgG into colostrum (3.9-fold transfer for total IgG, 11-fold transfer for bovine respiratory syncytial virus-specific IgG) (34). Together this suggests that virus-specific IgG is favored for transfer.

The mechanism required for transfer of IgG into canine colostrum is unclear. This makes understanding selective transfer of virus-specific IgG into colostrum even more challenging. The neonatal Fc-receptor (FcRn) has been implicated in IgG transfer into colostrum in some species (cattle, mice and pigs), however conflicting results indicate more research is still required (18,35). A role for FcRn in canine IgG transport into colostrum has been inferred based on studies in different species, but direct evidence is lacking. Identification of preferential virus-specific IgG transfer in the canine mammary gland supports the theory that this process could be receptor-mediated, and we suggest that features of virus-specific IgG are boosting this transfer. This process could be influenced by various characteristics of the IgG molecule which can affect its binding ability to FcRn, for example IgG subclass (20) or post translational modifications such as glycosylation of the Fc region. In cows, the most abundant IgG subclass found in colostrum is IgG1 (36), which is influenced by FcRn binding affinity, but other factors are also involved. We did perform preliminary investigations to evaluate the role of IgG subclasses in this study, but did not identify any change in transfer ratio for IgG2 compared to all IgG subclasses. We acknowledge our studies were limited by the lack of reagents available to precisely characterize the four canine IgG subclasses A-D (20,21). Regarding a potential role for antibody glycans in influencing MatAb transfer, evidence for glycosylation influencing transfer into colostrum comes from a recent study demonstrating that bovine colostrum contains IgG with higher levels of sialic acid compared to pooled serum (34). Moreover, vaccination in humans has been shown to induce different glycans in virus-specific antibodies but not all IgG (37). Future work is required to further understand this process in different species.

In this study, we identified maternal serum IgG titer as the strongest factor influencing the MatAb titers delivered to puppies. This finding is in contrast to a study by Mila et al, who studied 44 dogs in a single breeding kennel and did not identify an association between maternal serum total IgG titer and colostrum IgG titer (7). Interestingly, this earlier study reported a 2.8-fold enrichment of total IgG in colostrum, which is comparable to the 3.2-fold increase in total IgG we observed in our cohort. We suggest that the small difference in total IgG, and the lack of correlation between maternal IgG titer and colostrum titer in the earlier study is influenced by sampling strategy and timing; our colostrum samples were collected within one hour of delivery, whereas Mila et al. obtained samples up to 24 hours postpartum. Colostrum composition is known to change rapidly as it transitions to mature milk; in cattle, IgG concentration one day post-calving decreases to about 60% of the level measured at 0.5 days (34). This rapid decline coincides with intestinal ‘closure’, when neonates lose the ability to absorb macromolecules such as IgG. Gut closure occurs within 24 hours in both dogs and cows (Baker *et al.*, 2025), so consequently even small differences in sampling time may substantially influence colostrum IgG concentrations across studies. In addition, we observed a weaker association between total IgG titers in serum and colostrum than the correlation we identified between virus-specific IgG in colostrum and maternal serum, so this could contribute to the discrepancy in our findings. When considering other species, there is evidence that maternal serum IgG correlates with colostrum IgG in cats (38). Felidae have a similar endotheliochorial placenta to dogs, so similarities in MatAb transfer may be expected.

The evidence is less clear for species with epitheliochorial placentas however; a study in horses showed no correlation between maternal serum and colostrum IgG (39), and bovine studies have generated mixed results.

We did not detect any correlation between IgG titers in colostrum and parity, similar to findings in pigs (40). Interestingly, in cows, parity has been shown to modulate colostral IgG, with higher IgG concentrations observed in dams with increasing parity (41). These differences may reflect distinct reproductive and breeding practices among species, including the age at first breeding, which can influence colostrum composition across successive parturitions. Dam age did not appear to influence IgG titers in colostrum or maternal serum, and as expected, dam body weight did not correlate with IgG titers in colostrum or cord, consistent with prior findings that breed or size does not affect colostral IgG composition (1). In this cohort we also did not detect any association between reported vaccination status and MatAb titers, but recruitment of dogs during emergency surgery limited the completeness of our vaccination dataset.

Our discovery that colostrum and cord IgG titers correlate with dam IgG titers provides strong evidence that the most efficient way to boost protection in pups will be to boost antibody titers in the dam. We have shown a proportion of pups are born with protective titers of CPV-specific antibodies prior to suckling, and our study suggests this proportion could be increased by ensuring higher titers in dams. In our population however, 15.9% of the dams lacked detectable IgG titers to at least one highly pathogenic virus, leaving their puppies completely unprotected despite colostrum intake. As vaccinating dogs during pregnancy is currently not recommended by the World Small Animal Veterinary Association vaccine guidelines (16), it is essential that vaccine status prior to breeding is evaluated. We recommend administering booster vaccinations before breeding if the dam’s vaccination status is uncertain. Ensuring adequate anti-viral IgG titers in the dam will support effective placental transfer and promote the production of IgG-rich colostrum to maximize the delivery of protective MatAbs to puppies.

## ACKNOWLEGMENTS

We gratefully acknowledge the Riney Canine Health Centre for funding this project. The authors also express their gratitude for the Michael J. Day Scholarship to Lotta H Truyen. This scholarship is administered by the World Small Animal Veterinary Association and generously funded by MSD Animal Health. We also wish to thank Wendy Weichart for VLP production support, all owners of dogs recruited to this study, and the veterinary teams, with a special mention to Quinn Harting at Cornell University Hospital for Animals.

