## Supplementary figures and images for "Quantifying maternal antibody transfer to colostrum and cord blood reveals virus-specific selectivity in dogs"

### Supplemental Figure 1

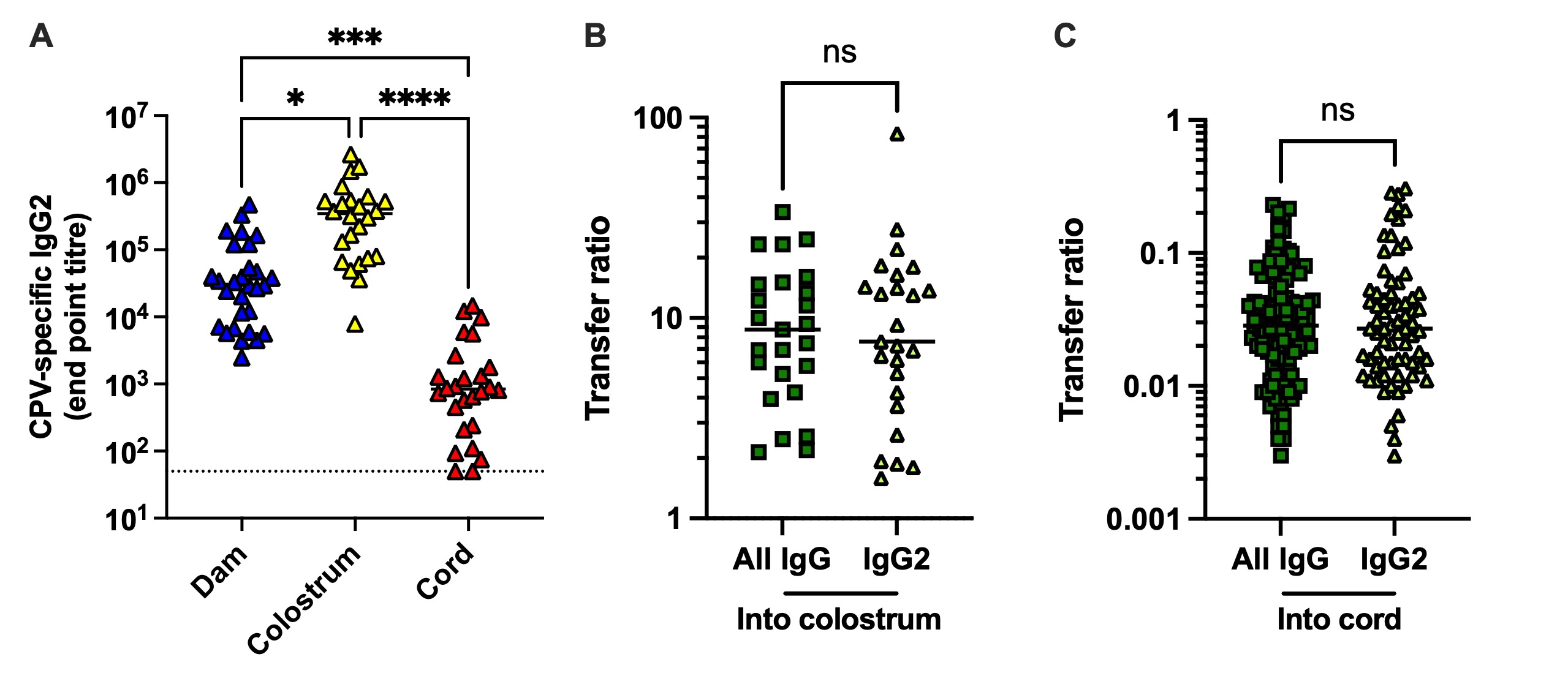

### Supplemental Figure 2

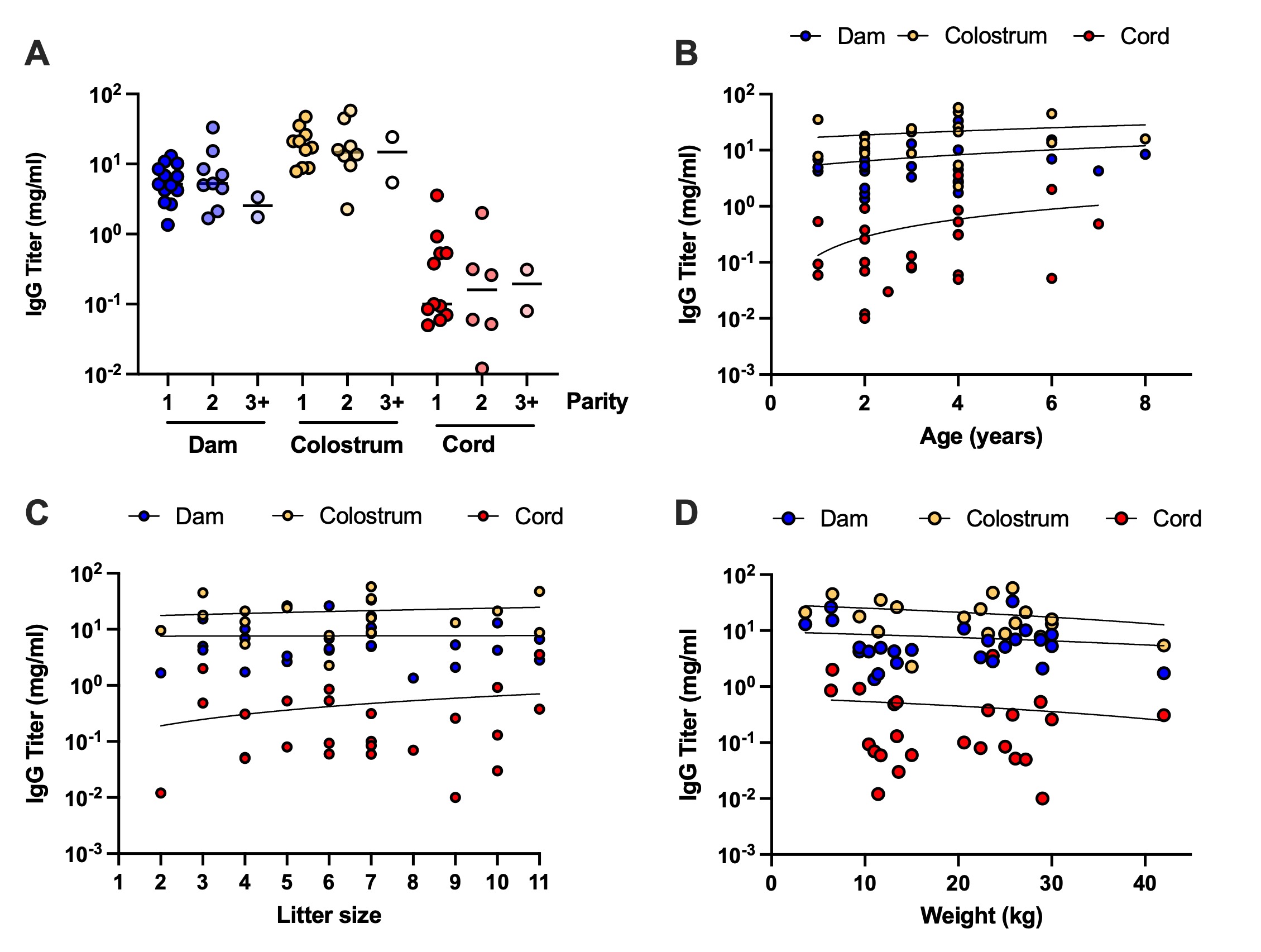
