## Supplemental Figure legends for "Quantifying maternal antibody transfer to colostrum and cord blood reveals virus-specific selectivity in dogs"

**Supplementary Figure Legends**

**Supplementary Figure 1.** **CPV-specific IgG2 quantified in dam serum, colostrum and cord samples.** A) End point titers of CPV-IgG2 specific MatAbs were quantified in clinical samples; dam n=29, colostrum n=24 and cord n=26 litters. Transfer ratios between dam serum and colostrum (B) and dam serum and cord serum (C) are shown. Dotted line in A represents lower limit of quantification. Significance was determined using Kruskal Wallis tests with multiple comparisons (A), and Wilcoxon tests (B/C). Asterisks denote statistical significance: *p* <0.01 (**)*, p* <0.001 (***), and *p* <0.0001 (****).

**Supplementary Figure 2. Correlation between total IgG titers and biological variables**. All values were determined by total IgG ELISA. (A) Total IgG (mg/ml) from dam, colostrum and cord samples plotted against parity. Horizontal bar represents the mean titer of each group. (B-D) Total IgG plotted against dam age, dam weight and litter size respectively. Non-linear regression curves were fitted to the data.
